# Towards ecologically valid biomarkers: real-life gait assessment in cerebellar ataxia

**DOI:** 10.1101/802918

**Authors:** Winfried Ilg, Jens Seemann, Martin Giese, Andreas Traschütz, Ludger Schöls, Dagmar Timmann, Matthis Synofzik

**Author notes:** Corresponding author: Winfried Ilg, Department of Cognitive Neurology, Hertie Institute for Clinical Brain Research, Otfried-Müller-Straße 25, 72076 Tübingen, Germany.

## Abstract

**BACKGROUND:** With disease-modifying drugs on the horizon for degenerative ataxias, motor biomarkers are highly warranted. While ataxic gait and its treatment-induced improvements can be captured in laboratory-based assessments, quantitative markers of ataxic gait in real life will help to determine ecologically meaningful improvements.

**OBJECTIVES:** To unravel and validate markers of ataxic gait in real life by using wearable sensors.

**METHODS:** We assessed gait characteristics of 43 patients with degenerative cerebellar disease (SARA:9.4±3.9) compared to 35 controls by 3 body-worn inertial sensors in three conditions: (1) laboratory-based walking; (2) supervised free walking; (3) real-life walking during everyday living (subgroup n=21). Movement analysis focussed on measures of movement smoothness and spatio-temporal step variability.

**RESULTS:** A set of gait variability measures was identified which allowed to consistently identify ataxic gait changes in all three conditions. Lateral step deviation and a compound measure of step length categorized patients against controls in real life with a discrimination accuracy of 0.86. Both were highly correlated with clinical ataxia severity (effect size ρ=0.76). These measures allowed detecting group differences even for patients who differed only 1 point in the SARA_p&g_ subscore, with highest effect sizes for real-life walking (*d*=0.67).

**CONCLUSIONS:** We identified measures of ataxic gait that allowed not only to capture the gait variability inherent in ataxic gait in real life, but also demonstrate high sensitivity to small differences in disease severity - with highest effect sizes in real-life walking. They thus represent promising candidates for quantitative motor markers for natural history and treatment trials in ecologically valid contexts.

## Introduction

Gait disturbances often present as the first signs of degenerative cerebellar ataxia (DCA) ^1, 2^ and are one of the most disabling features throughout the disease course. It has been shown in laboratory-based assessments that measures of spatial and temporal movement variability allow distinctively to capture and characterize the specificities of ataxic gait ^3-10^. Moreover, they allow to quantify disease severity even at preclinical stages of DCA ^11, 12^ and to capture treatment-induced improvements in a fine-grained fashion ^13-15^, thus suggesting a high potential as both progression and treatment response parameters in the upcoming treatment trials ^16-18^. Recently, first studies showed that such measures of spatio-temporal variability characterizing ataxic gait can also be captured using wearable inertial sensors in clinical assessments ^19, 20^. With their easy handling outside from specialized gait laboratories, these systems are particularly promising for conduction of multi-centered studies. Moreover, driven by advances in sensor technologies, wearable sensors increasingly enable remote capturing of patients’ movements in their real life. This might allow to capture patients’ real-world motor performance and ecologically meaningful intervention-induced improvements, rather than “snapshot” performance during constrained and partly “artificial” surrogate motor tasks assessed by clinical scores or lab conditions ^21^.

While wearable sensors have already proven their value to capture and quantify characteristics of real-life movement in neurological diseases like Parkinson’s disease ^22, 23^ or multiple sclerosis ^24^, studies are still lacking which capture the specifics of ataxic gait in real life beyond the level of mere physical activity ^25-27^.

The transfer of spatio-temporal variability measures for quantifying ataxic gait impairments into real life is hereby complicated by the fact that real life gait is in general inherently far more variable for both healthy controls and cerebellar patients ^28^ and that patients are free to use various compensation strategies, thus increasing heterogeneity of walking patterns and complexity of gait scenarios. Thus, spatio-temporal gait measures may lose their accuracy for characterizing ataxic gait changes in real life.

Here we aimed to unravel spatio-temporal gait measures in real-life environments that allow to quantify not just unspecific walking changes in ataxia (e.g. decreased walking speed), but features inherent to ataxic gait changes in DCA-as it will be these features that will be particularly sensitive to change by upcoming treatment trials specifically targeting cerebellar dysfunction.

## Methods

### Participants

43 patients with degenerative cerebellar ataxia (CA, age: 51 ±15 years) were recruited from the Ataxia Clinics of the University Hospitals Tübingen and Essen. Patients were included based on following inclusion criteria: 1.) degenerative cerebellar ataxia in the absence of any signs of secondary CNS disease; 2.) age between 18 and 75 years; 3.) able to walk without walking aids. The exclusion criteria were: severe visual or hearing disturbances, cognitive impairment, predominant non-ataxia movement disorders (e.g. parkinsonism, spasticity), or orthopaedic constraints. Severity of ataxia was rated using the SARA (Scale for the Assessment and Rating of Ataxia) ^29^. SARA covers a range from 0 (no ataxia) to 40 (most severe ataxia). The SARA score includes the following 8 items: 3 items rating gait and posture, 1 item for speech disturbances, and 4 items for limb-kinetic functions. The 3 items rating gait and posture are grouped by the subscore SARA posture & gait (SARA_p&g_) ^11, 30^. The group of cerebellar patients had a mean SARA score of 9.4 (range [1-16]) and mean SARA_p&g_ subscore of 3.6 (range [0-6]). In addition, we recruited 35 healthy controls (HC, age: 48±15 years). HC subjects had no history of any neurological or psychiatric disease, no family history of neurodegenerative disease, and did not show any neurological signs upon clinical examination. The experimental procedure was approved by the local ethics committee (598/2011BO1). All participants gave their informed consent prior to participation.

### Gait Conditions

Walking movements were recorded in three different conditions, namely: (i) **L**ab-**b**ased **w**alking (LBW condition): walking was constrained by a specified walking distance of 50m in a specific quiet non-public indoor floor within an institutional setting (hospital), and supervised by a study assessor watching the walking performance; patients were instructed to walk normally at a self-selected speed; (ii) **S**upervised **F**ree **W**alking (SFW condition): unconstrained walking in public indoor floor and outdoor spaces on an institutional (hospital) compound where subjects were free to choose and change the floors and indoor and outdoor spaces where they wished to walk (complete walking time: 5 minutes) with all spaces being open to public, but still supervised by a study assessor watching subject’s walking performance; (iii) **R**eal-**L**ife **W**alking (RLW condition): unconstrained walking during subjects’ everday living where subjects where free to move how they wanted and were used to in their individual daily life, without supervision by any study personnel (total recording time: 4-6 hours). Subjects were instructed to wear the sensors inside and outside their house, and include at least a half-hour walk (for an overview of all conditions, see Supplement S1 Table 1). Subjects documented their recorded walking movements in an activity protocol. Out of the respective total groups, 21 CA patients and 17 HC were available for the RLW condition (for an overview of these subjects, see Table 1).

**Table 1.**
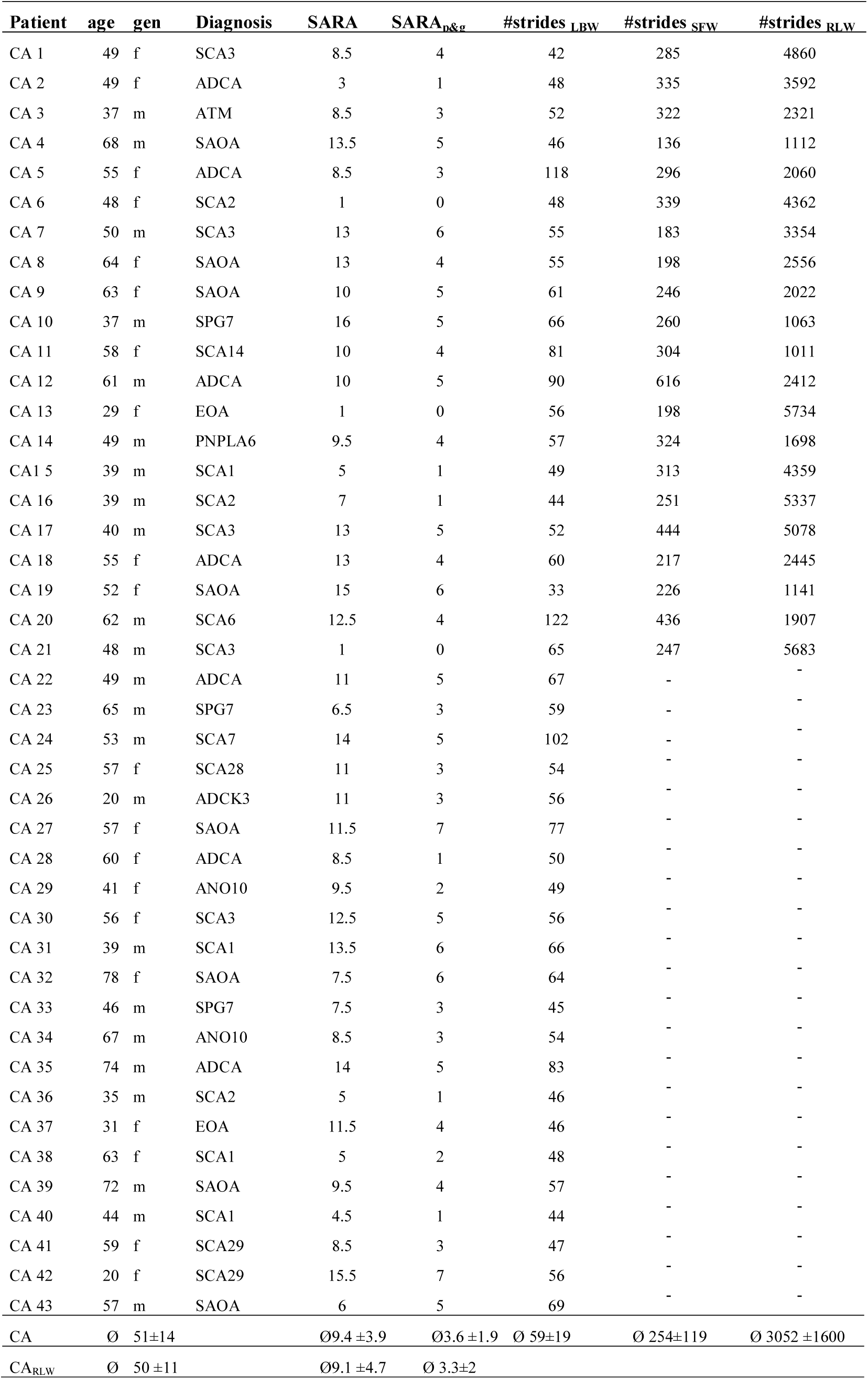
Patient characteristics. Clinical ataxia severity was determined by the SARA ^29^. The SARA subscore posture&gait is defined by the first three items of the SARA score which capture gait, standing and sitting ^30^. #Strides denotes the number of steps analysed for the given walking condition. SAOA: sporadic adult onset ataxia; ADCA: autosomal dominant ataxia of still undefined genetic cause; EOA: early-onset ataxia of still undefined genetic cause. SCA: autosomal-dominant spinocerebellar ataxia of defined genetic type. The following diagnosis denote the gene underlying the respective ataxia type: ATM (=Ataxia teleangiectasia), SPG7 (=hereditary spastic paraplegia type 7), SYNE1 (= Autosomal recessive cerebellar ataxia type I), SETX (= AOA2: Ataxia with oculomotor apraxia type 2), ADCK3 (=ARCA 2, Autosomal-recessive cerebellar ataxia type 2); PNPLA6, ANO10 (=SCAR 10, Autosomal-recessive spinocerebellar ataxia type 10).

### Movement measures

Three Opal inertial sensors (APDM, Inc., Portland, US) were attached on both feet, and posterior trunk at the level of L5 with elastic Velcro bands. Inertial sensor data was collected and wirelessly streamed to a laptop for automatic generation of gait and balance metrics by Mobility Lab software (APDM, Inc., Portland, US). For the unconstrained walking conditions (SFW, RLW), data was logged on board of each OPAL sensor and downloaded after the session. Selected walking bouts contained 5 subsequent strides with a minimum average velocity of 0.5 * the average walking speed in the constrained walking trail. Step events, as well as spatio-temporal gait parameters from the IMU sensors were extracted using APDM’s mobility lab software (Version 2)^31^, which has been shown to deliver good-to-excellent accuracy and repeatability ^32, 33^. For each detected stride, the following features were extracted: stride length, stride time, lateral step deviation and raw accelerometer data of the lumbar sensor. Variability measures were calculated using the coefficient of variation CV=σ/μ, normalizing the standard deviation with the mean value ^34^. On this basis, stride length CV (StrideL_CV_) and stride time CV (StrideT_CV_) were determined, which have shown to be sensitive to ataxia severity in constrained lab-based walking ^3, 5, 7, 11, 12^.

The measure of lateral step deviation (*LatStepDev*) was determined on the basis of three consecutive walking steps, calculating the perpendicular deviation of the middle foot placement from the line connecting the first and the third step (see Supplement S1, Figure 1). *LatStepDev* was normalized with stride length (% of stride length), thus providing a measure independent from step length variability, which is suggested to be increased in real-life gait.

In order to establish a measure which captures different types of spatial step variability, we combined the measure of step length variability (*StrideL*_*CV*_) (mostly anterior-posterior direction) with the measure of lateral step deviation (*LatStepDev*) (medio-lateral dimension). The spatial step variability compound measure *SPcmp* determines for each of the two parameters (*StrideL*_*CV*_) and (*LatStepDev*) the relative value of an individual subject in comparison to the value range of all subjects (resulting in values between [0-1]) and takes the maximum of both values (see also Figure 2 in Supplement S1).

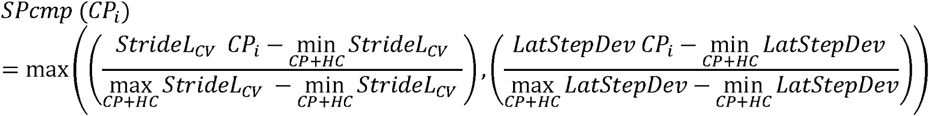

In addition, harmonic ratio (HR) ^35, 36^ of pelvis acceleration was determined to quantify the smoothness of motion. The method quantifies the harmonic content of the acceleration signals in each direction (HR AP: anterior-posterior, ML medio-lateral, V: vertical) using stride frequency as the fundamental frequency component. Using a finite Fourier series, the components of the acceleration signal that are ‘in phase’ (the even harmonics) are compared to the components that are ‘out of phase’ (odd harmonics), and a harmonic ratio is calculated by dividing the sum of the amplitudes of the first ten even harmonics by the sum of the amplitudes of the first ten odd harmonics 35. Thus, the HR quantifies the harmonic composition of these accelerations for a given stride where a higher HR is interpreted as greater walking smoothness (see Supplement S1 for a more formal description). It has been recently shown that HR measures distinguish between cerebellar patients and healthy controls in lab-based walking trails ^20, 37^

### Statistics

Between-group differences (CA vs. HC group) of movement features were determined by the non-parametric Kruskal-Wallis-test. When the Kruskal-Wallis-test yielded a significant effect (p<0.05), post-hoc analysis was performed using a Mann-Whitney U-test. The same tests were used to distinguish between ataxia severity subgroups, which were stratified according to the degree of gait and posture dysfunction based on the SARA_p&g_ subscore: mild CA_Mild_ =SARA_g&p_ [0-2]; moderate CA_Mod_: =SARA_g&p_ [3-4]; severe CA_Sev_:=SARA_g&p_ [5-6]. Receiver operating characteristic (ROC) analysis determining the classification accuracy was used to quantify the discrimination capability of the examined measures for different walking conditions.

Repeated measurements analyses were performed using the non-parametric Friedman test (χ^2^,p-values) to determine within-group differences between walking conditions. When the Friedman test yielded a significant effect (p<0.05), post hoc analysis was performed using a Wilcoxon signed-rank test for pairwise comparisons between assessments. We report three significance levels: (i) uncorrected *:p<0.05, (ii) Bonferroni-corrected for multiple comparisons **: p<0.05/n=7: number of analysed features, (iii) ***p<0.001. Spearman’s ρ was used to examine the correlation between movement measures and SARA scores as well as between measures for different walking conditions. Effect sizes were classified as ρ: 0.1 small effect, 0.3 medium effect, 0.5 large effect, 0.7 very large effect ^38, 39^.

To analyse the sensitivity of the movement measures to detect clinically important changes in ataxia severity, we compared gait measures for patients with (i) a difference of 1 point as well as (ii) with a difference of 2 points in the SARA_p&g_ subscore. These ranges are motivated by previous analysis on the responsiveness of the SARA, showing that a change of 1 SARA point can be considered as a clinically important progression ^40^. In addition, they are motivated by motor intervention studies demonstrating that current treatment interventions can yield an average improvement of 1.5-2 points on the SARA_p&g_ subscore, and that these effects represent patient-relevant improvements ^14, 15, 41, 42^. A Wilcoxon signed-rank test was performed to test for differences between patient groups categorized by a delta of 1 and 2 points SARA_p&g_, respectively. The effect sizes for group differences were determined by Cohen’s *d* ^38^ with pooled standard deviations ^43^ (cohen’s *d* = [0.5-0.8]: medium effect; d> 0.8: strong effect; d> 1.3: very strong effect ^38^). Statistical analysis was performed using MATLAB (Version2017 B).

## Results

### Group differences between HC and CA for different walking conditions

In the constrained walking condition (LBW, N_CA_=43), highly significant group differences (p< 0.00014) were observed for all measures of spatio-temporal gait variability and smoothness of movements (Figure 1; similar LBW results also if considering only that subgroup of CA patients who were available also for the RLW condition [LBW_RLW-subgroup,_ n=21], see Supplement S2, Table 2). Also in the unconstrained walking conditions SFW and RLW, several variability measures like *StrideL*_*CV*_ (p=0.025, *d* =0.82, *ROC accuracy 0.75*) and *StrideT*_*CV*_ (p=0.02, *d* =0.86, *ROC accuracy 0.72*) allowed to distinguish between HC and CA (see Figure 1 and Supplement S2, Table 2). Effect sizes and discrimination performance were smaller than in LBW, as variability in gait measures was generally higher in these unconstrained walking conditions, which was observed in both healthy controls and patients.

**Figure 1.**
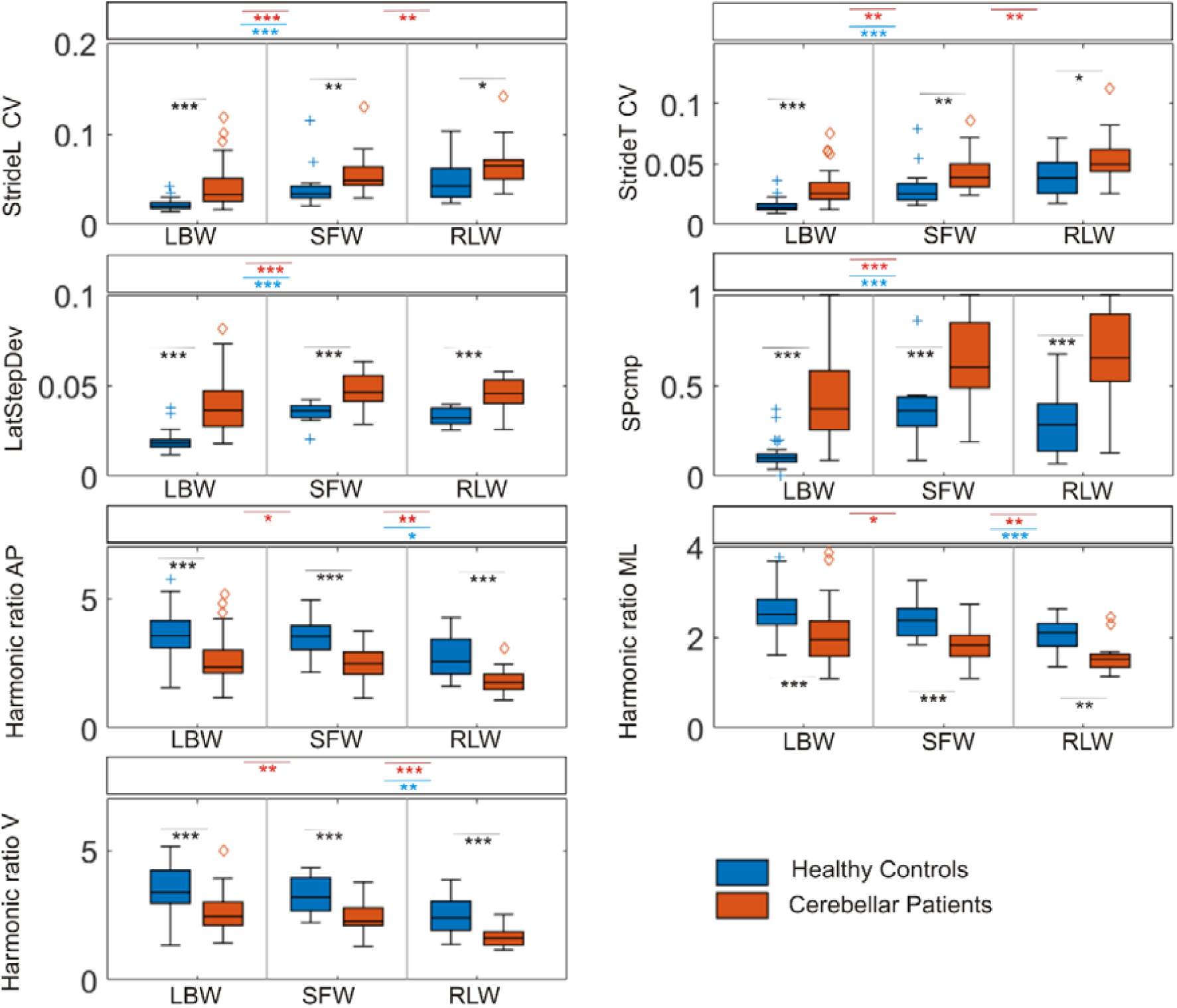
Shown are *between-group differences* between cerebellar patients (CA, orange) and healthy controls (HC, blue) *within* each of the different walking conditions. Shown are group differences for constrained lab-based walking (LBW), supervised free walking (SFW) and real life walking (RLW). Black stars indicate significant differences between groups (*= p<0.05, **= p<0.007 Bonferroni-corrected, ***= p<0.001). Shown are also within-group differences *between* the different walking conditions (orange stars: significant differences between walking conditions in the CA cohort; blue stars: significant differences between walking conditions in HC).

In contrast, the measures *LatStepDev* and *SPcmp* showed a similarly high effect size for both unconstrained conditions (SFW, RLW) as in the constrained condition LBW, which was observed in both patients and controls (Figure 1, see blue and red stars for differences between conditions). These measures also showed the clearest discrimination between CA and HC in the real-life condition RLW *(LatStepDev*: p=0.0002, *d=1.6, ROC accuracy 0.86*; *SPcmp: p=0.00012, d=2.6, ROC accuracy 0.86)*.

### Sensitivity to ataxia severity in different walking conditions

Most movement measures in the constrained walking condition LBW showed a highly significant correlation with the SARA_p&g_ subscore, (effect size ρ>*0.65*), indicating a sensitivity of our measures in this condition to ataxia severity (Table 2). The degree of these correlations decreased for several measures in the unconstrained walking condition SFW, and for even more measures in the real-life condition RLW, including *StrideL*_*CV*_ and *StrideT*_*CV*_. However, the measures *LatStepDev, SPcmp* and *Harmonic Ratios (AP)* revealed significant correlations of high effect size with the SARA_p&g_ subscore (p≤0.008, ρ > 0.56) also in the condition RLW (Table 2).

**Table 2.** Correlations between the SARA_p&g_ subscore and gait measures in different walking conditions for the cohort of cerebellar patients (*= p<0.05, **= p<0.007 Bonferroni-corrected, ***= p<0.001). Effect sizes of correlations are given using Spearman’s ρ.

| Correlation<br>SARA<sub>p&g</sub>/<br>gait measures | constrained walking<br>LBW |  | free walking<br>SFW |  | real life walking<br>RLW |  |
| --- | --- | --- | --- | --- | --- | --- |
|  | ρ | p | ρ | p | ρ | p |
| StrideL<sub>CV</sub> | 0.65 | <0.0001*** | 0.34 | 0.1 | 0.47 | 0.03* |
| StrideT<sub>CV</sub> | 0.71 | <0.0001*** | 0.2 | 0.36 | 0.27 | 0.24 |
| LatStepDev | 0.75 | <0.0001*** | 0.63 | 0.001** | 0.63 | 0.0023** |
| SPcmp | 0.78 | <0.0001*** | 0.64 | <0.0001*** | 0.76 | 0.0001*** |
| Harmonic Ratio AP | -0.58 | 0.0004*** | -0.6 | 0.002** | -0.56 | 0.008* |
| Harmonic Ratio ML | -0.34 | 0.026* | -0.56 | 0.0049** | -0.47 | 0.033* |
| Harmonic Ratio V | -0.55 | 0.00011*** | -0.55 | 0.0051** | -0.47 | 0.033* |

In order to examine the sensitivity of the movement measures to ataxia severity in further detail, we binned the patient population in three subgroups: mild *CA*_*Mild*_ (#6 subjects in RLW), moderate *CA*_*Mod*_ (#7 subjects in RLW) and severe *CA*_*Sev*_ (#8 subjects in RLW) according to the SARA_p&g_ subscore (see methods). Figure 2 shows subgroup measures for the walking conditions LBW and RLW. The measures StrideT_CV_ (p<0.006**), LatStepDev (p<0.02*), SPcmp (p<0.01*) and HR AP (p<0.01*) distinguished significantly between subgroups for constrained walking (LBW). Moreover, SPcmp distinguished between subgroups in real life (p<0.03*), despite the small sizes of the subgroups in the RLW condition.

**Figure 2.**
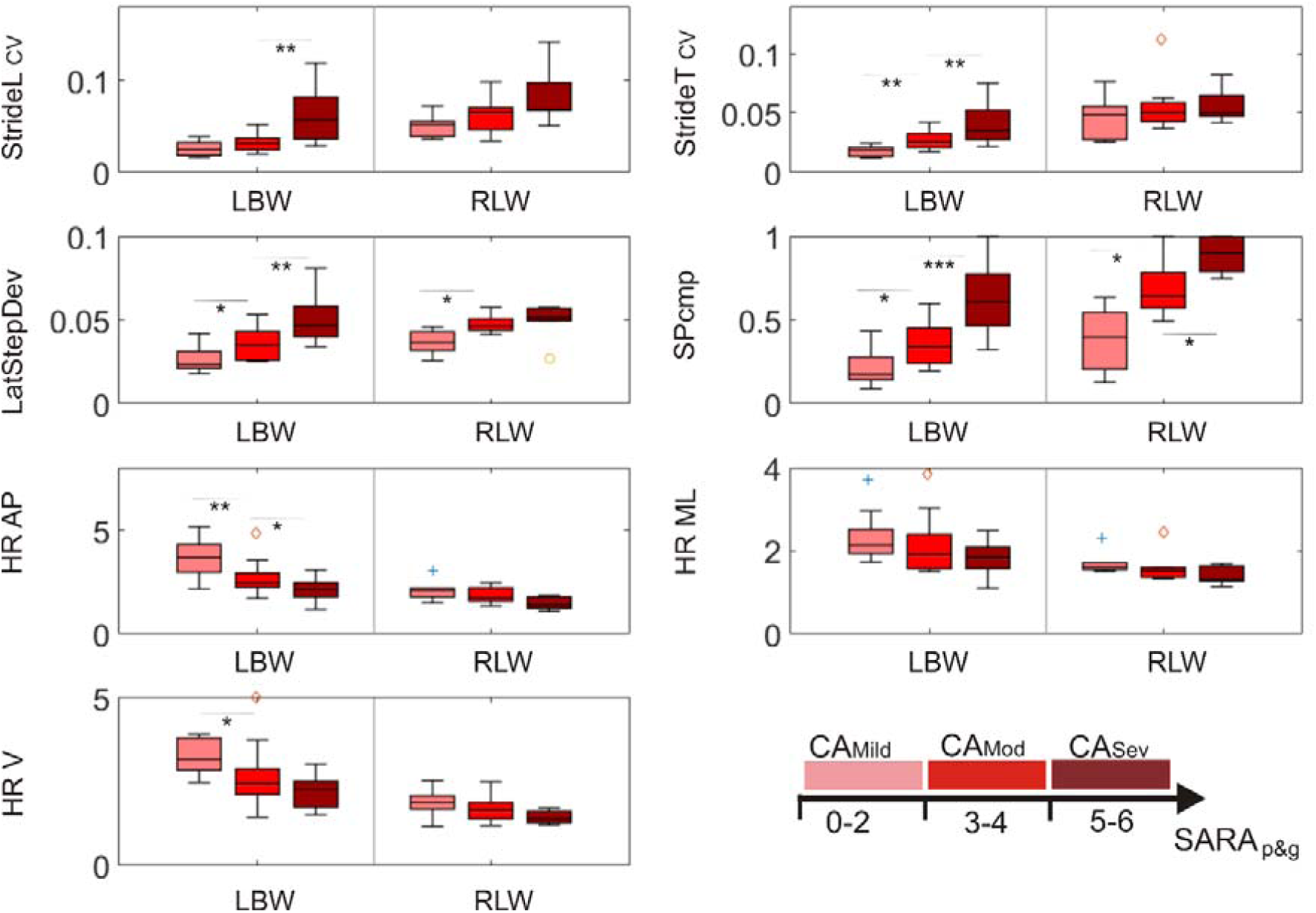
Differences between subgroups of cerebellar patients stratified according to gait and posture ataxia severity as determined by the SARA_p&g_ subscore. Subgroups: CA_Mild_: SARA_g&p_ [0:2], CA_Mod_: SARA_g&p_= [3-4], CA_Sev_: SARA_g&p_ [5 - 6]. Shown are group differences for constrained lab-based walking (LBW) and real life walking (RLW).

### Sensitivity to capture clinically important differences in ataxia severity in real life

We next analysed whether our measures allow to detect the quantitative motor correlates of rather small but clinically important and everyday-living relevant differences (see methods and discussion). To this end, we compared measures for patients who differ only 1 and 2 points, respectively, in the SARA_p&g_ subscore. Paired statistics revealed significant differences between these patient groups for several measures (Table 3). The compound measure *SPcmp* yielded the largest effect sizes for the real-life condition (RLW) of *d* =0.67 for Δ SARA_p&g_=1, and *d* =1.2 for Δ SARA_p&g_=2. Despite smaller cohort size (N_LBW_=43, N_RLW_=21), effects sizes in the RLW condition outperform those of the LBW condition (Table 3).

**Table 3.** Differences between gait measures (p-values, *Wilcoxon signed-rank test* plus effect sizes indicated by Cohen’s *d*) when patients differ in SARA_p&g_ subscore by one (Δ SARA_p&g_=1) or two points (Δ SARA_p&g_=2), respectively. Shown are results from all walking conditions LBW, SFW and RLW as well as for LBW with the subgroup of patients who were also available for the RLW condition (LBW_RLW-subgroup_).

| Walking condition | Measure | p ΔSARA<sub>p&g</sub>=1 | Cohen's *d* | p ΔSARA<sub>p&g</sub>=2 | Cohen's *d* |
| --- | --- | --- | --- | --- | --- |
| **LBW** | StrideL<sub>CV</sub> | <0.0001*** | 0.42 | <0.0001*** | 0.78 |
|  | StrideT<sub>CV</sub> | <0.0001*** | 0.38 | <0.0001*** | 0.79 |
|  | LatStepDev | <0.0001*** | 0.49 | <0.0001*** | 0.87 |
|  | SPcmp | <0.0001*** | 0.51 | <0.0001*** | 0.92 |
|  | HR AP | 0.0041** | 0.23 | <0.0001*** | 0.81 |
|  | HR ML | 0.088 | 0.15 | 0.0001*** | 0.4 |
|  | HR V | 0.011* | 0.22 | <0.0001*** | 0.6 |
| **LBW<sub>RLW-subgroup</sub>** | StrideL<sub>CV</sub> | 0.0047** | 0.51 | 0.0094* | 0.78 |
|  | StrideT<sub>CV</sub> | 0.023* | 0.38 | 0.015* | 0.67 |
|  | LatStepDev | 0.048* | 0.28 | 0.019* | 0.58 |
|  | SPcmp | 0.0087* | 0.39 | 0.007** | 0.74 |
|  | HR AP | 0.39 | 0.14 | 0.016* | 0.69 |
|  | HR ML | 0.081 | 0.33 | 0.81 | 0.06 |
|  | HR V | 0.27 | 0.16 | 0.21 | 0.37 |
| **SFW** | StrideL<sub>CV</sub> | 0.096 | 0.28 | 0.03* | 0.44 |
|  | StrideT<sub>CV</sub> | 0.53 | 0.1 | 0.18 | 0.29 |
|  | LatStepDev | <0.0001*** | 0.49 | 0.0093* | 0.61 |
|  | SPcmp | <0.0001*** | 0.51 | 0.0016** | 0.65 |
|  | HR AP | 0.14 | 0.22 | 0.0003*** | 0.79 |
|  | HR ML | 0.011* | 0.4 | 0.0021** | 0.59 |
|  | HR V | 0.072 | 0.28 | 0.0011** | 0.62 |
| **RLW** | StrideL<sub>CV</sub> | 0.02* | 0.42 | 0.09 | 0.52 |
|  | StrideT<sub>CV</sub> | 0.45 | 0.14 | 0.76 | 0.09 |
|  | LatStepDev | 0.053 | 0.33 | 0.043* | 0.59 |
|  | SPcmp | <0.0001*** | 0.67 | <0.0001*** | 1.2 |
|  | HR AP | 0.0007*** | 0.6 | 0.034* | 0.64 |
|  | HR ML | 0.0017** | 0.57 | 0.27 | 0.33 |
|  | HR V | 0.0007*** | 0.61 | 0.22 | 0.37 |

### Relationships of movement measures across conditions and across measures

All measures of spatial and temporal variability as well as the harmonic ratios were highly correlated across all three conditions in cerebellar patients (Supplement S2 Table 5). In contrast, only few correlations were found across conditions in healthy controls (Supplement S2 Table 6). Similarly, also close relationships across measures were observed for cerebellar patients, with strong correlations between harmonic ratios determining movement smoothness and spatio-temporal step variability for all walking conditions (Supplement S2, Table 7). They were more pronounced in cerebellar patients than in healthy controls (Supplement S2, Table 7).

## Discussion

This study aimed to test the hypothesis that spatio-temporal gait measures reflecting the inherent features of ataxic gait in DCA can be captured by wearable sensors not only in indoor and supervised clinical settings, but also remotely during real-life walking in everyday living. We were able to identify measures which allow to quantify ataxia features across all of these settings with high discrimination accuracy against controls as well as sensitivity to ataxia severity. This included in particular unconstrained real-life environments (RLW) as more complex, yet ecologically more valid settings, e.g. for future patient-centered treatment trials in DCA.

### Spatio-temporal gait variability of as a consistent feature of ataxic gait

Our findings in the constrained walking condition LBW confirm the results of previous studies from our and other groups with different movement capture technologies ^3, 4, 6, 12, 44^, including wearable sensors ^19^, ^45-47^. These studies showed that spatio-temporal variability measures like stride length variability (*StrideL*_*CV*_) and stride time variability (*StrideT*_*CV*_) in constrained walking serve as reliable and valid measures for cerebellar ataxia and - as demonstrated here for wearable sensors -correlate well with gait and posture ataxia severity. Moreover, first studies using wearables sensors have indicated that gait analysis might be more responsive for one year ataxia progression changes than the SARA score ^47^. Taken together with our current observations, these findings are important as they confirm that measures of spatio-temporal variability deliver consistent, reproducible results in ataxia patients across methods and centres, as warranted for upcoming multicentre natural history and treatment trials in DCA.

However, our findings add additional promising measures for ataxic gait, with *LatStepDev* and *SPcmp* showing higher effects sizes and discrimination accuracy of DCA against controls than the aforementioned previous measures (which was observed also for the constrained walking condition LBW). In addition, also harmonic ratios representing measures of trunk movement smoothness - initially used in Parkinson’s Disease ^48^ and Multiple Sclerosis ^49^ and more recently also in CA ^20, 37^ – show high sensitivity for ataxia severity in constrained movements, indicating their value as novel measures quantifying ataxic gait.

### From constrained gait to real life walking

We observed an increased *within-group* spatio-temporal variability of the measures *StrideL*_*CV*_ and *StrideT*_*CV*_ in both healthy controls and cerebellar patients in real-world walking (condition RLW) compared to supervised constrained walking in a clinical setting (condition LBW) (Figure 1). This observation, which is consistent with previous work confined to healthy subjects so far ^28^, can be explained by increased voluntary variation of step length in real-life gait behaviour.

This increased spatio-temporal variability in real life led to a decrease in effect size and discrimination accuracy for common measures of step variability like *StrideL*_*CV*_ and *StrideT*_*CV*_ in the group comparison of cerebellar patients compared to healthy controls (Supplement S2 Tables 3+4). Yet, large effect sizes and discrimination accuracies even in the real-life condition were revealed for the measure *LatStepDev* and the new compound measure *SPcmp,* with high similarity of these measures across conditions. This indicates that *LatStepDev* and *SPcmp* may capture a more condition independent, i.e. robust ataxia component of spatio-temporal variability than *StrideL*_*CV*_ and *StrideT*_*CV*_.

### Measures of ataxic gait during real life: sensitivity to ataxia severity

The measures *LatStepDev* and *SPcmp* as well as *harmonic ratios AP* and *V* did not only allow to distinguish cerebellar patients from healthy controls in real life; they were also highly correlated to clinical ataxia severity in this condition (see Table 2). While *harmonic ratios, StrideL_CV_*, and *StrideT*_*CV*_ failed to reach significance for differentiating the three severity subgroups of cerebellar patients for the real-life condition RLW, *LatStepDev* and *SPcmp* were sensitive to distinguish these severity subgroups also during real-life walking (Figure 2). The compound measure *SPcmp* hereby seems to benefit from capturing different compensation strategies used in diverse stages of disease, and might allow to capture gait ataxia in particular in more advanced disease stages (for a more detailed description on the characteristics of the measure *SPcmp*, see Supplement S1 Figure 3).

### Sensitivity to capture important differences in real life for future natural history and intervention trials

To serve as progression and treatment outcome measures, measures of real-life walking should ideally be able to capture changes that correspond to clinically and everyday-living important differences as well as to treatment effects achievable by current and future ataxia treatment interventions. A change of one point in the SARA score has been shown to reflect a clinically important difference over one year disease course ^40^, determined by the patients’ global impression of change (PGI) using quality of life outcomes ^50^. Moreover, changes of 1.5-2 points in the SARA_p&g_ subscore reflect treatment effect sizes consistently achieved by currently available motor rehabilitation interventions ^14, 15, 41, 42^. Our measure *SPcmp* yields a strong effect size (Cohen’s *d*=1.2) for differentiating movement patterns when patients differ by two SARA_p&g_ points, and an at least moderate effect size (Cohen’s d=0.67) when patients differ by one SARA_p&g_ point, demonstrating that this measure is able to capture clinically important differences. Remarkably, for both types of clinical important differences (Δ SARA_p&g_=1; Δ SARA_p&g_=2) the highest effect sizes were observed in the real-life condition RLW (Table 3), despite the general increase of variability in real time walking. This observation might be explained by the larger amount of walking strides available for analysis in this condition and, in addition, by the particular movement characteristics of unconstrained walking. In contrast, a shorter unconstrained trial – like the condition SFW which comprised of only 5 minutes walking – does not seem to yield equally large effect sizes. This observation is important as outcome measures with higher effect sizes -as observed here for the real-life walking condition - may reduce the sample sizes required in natural history studies and upcoming treatment trials in hereditary ataxias using e.g. antisense oligonucleotides ^16-18^.

### Relationships between clinical gait assessments and real-life walking

Despite the general increase of variability in real-life walking, we observed high correlations between the constrained lab-based (LBW) and the unconstrained (SFW, RLW) walking conditions. This suggests that also the lab-based assessment might be exploited to deliver first surrogate snapshots of patients’ unconstrained gait performance. However, as noted above, at least some of the measures seemed to yield larger effect sizes in real-life walking. Moreover, our current analysis of real-life walking behaviour was limited to walking bouts of minimal five subsequent strides (rather than analysis of more complex everyday-living walking behaviours) which might explain the good correlations with the constrained walking conditions. However, real life includes a much larger variety of walking movements, for instance turning movements or initiation and termination of gait, all known to be demanding for dynamic balance control and impaired in cerebellar disease. To include these movements in future analysis of real-life walking is highly warranted to capture ecological validity in even more depth.

## Conclusion and outlook

This study unravels measures that allow to quantify real-life ataxic gait and hereby reflect disease severity, thus yielding promising ecologically valid outcome measure candidates for future natural history and treatment trials in degenerative cerebellar ataxias. For both types of trials, measuring real-life movements bears several other advantages - in addition to the higher effect sizes gained from real-life assessments, likely caused by larger amount of sampled walking strides. These advantages include objective quantitative measurement of (i) day-to day variability instead of snapshot evaluations weeks or months apart during clinical visits ^21^ and of (ii) patients’ real-world motor performance instead of partly “artificial” motor tasks of clinical scores or lab conditions, which serve as surrogate parameters at best. While assessment of constrained tasks at snapshot visits represents patients’ real-world functioning only in a limited fashion ^51^, measures of real-life motor performance add ecological validity and can thus help to inform upcoming treatment trials in degenerative cerebellar ataxias and FDA approval of novel treatments.

## Acknowledgements

This project received support, in part, by the German Hereditary Ataxia Society (DHAG), the “Stiftung Hoffnung” (to M.S.) and the Clinician Scientist Programme of the University of Tübingen (439-0-0, to A.T.). Additional support has been received from BW Stiftung (project KONSENS NEU007/1 to M.G.).

## Supplement S1 – Details of gait conditions and measures

### Walking conditions

**Table 1.**
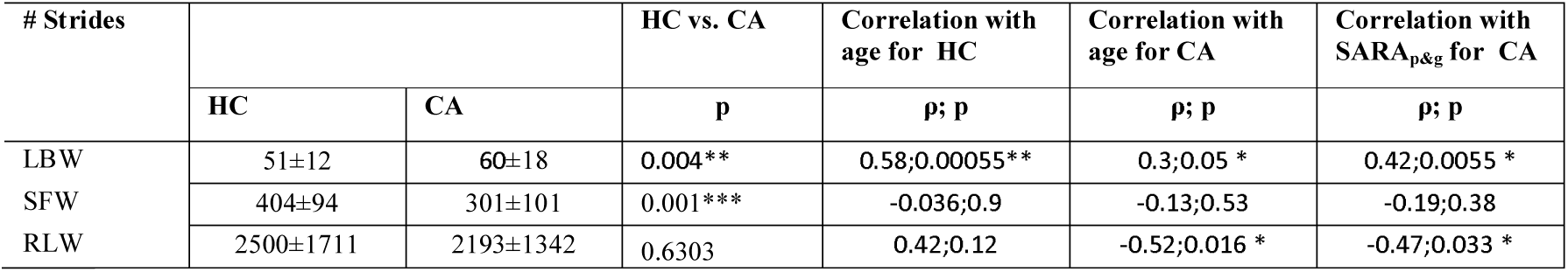
Description of walking conditions.

### Lateral Step Deviation

The measure of lateral step deviation (*LatStepDev*) was determined on the basis of three consecutive walking steps, calculating the perpendicular deviation of the middle foot placement from the line connecting the first and the third step (see Figure 1).

**Figure 1.**
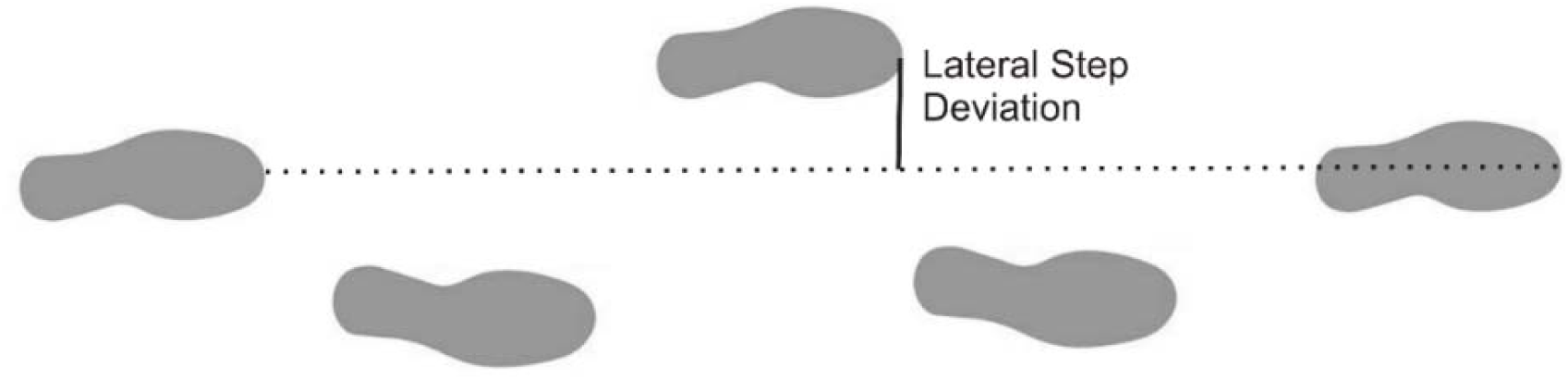
Illustration on the measurement of lateral step deviation (LatStepDev).

### Compound measure SPcmp

In order to establish a compound measure for spatial step variability *SPcmp*, we combined the measure of step length variability (StrideL_CV_) with the measure of lateral step deviation (LatStepDev). *SPcmp* determines for each of the two parameters (StrideL_CV_) and (LatStepDev) the relative value of an individual subject in comparison to all other subjects (resulting in values between [0-1]) and takes the maximum of both measures.

**Figure 2.**
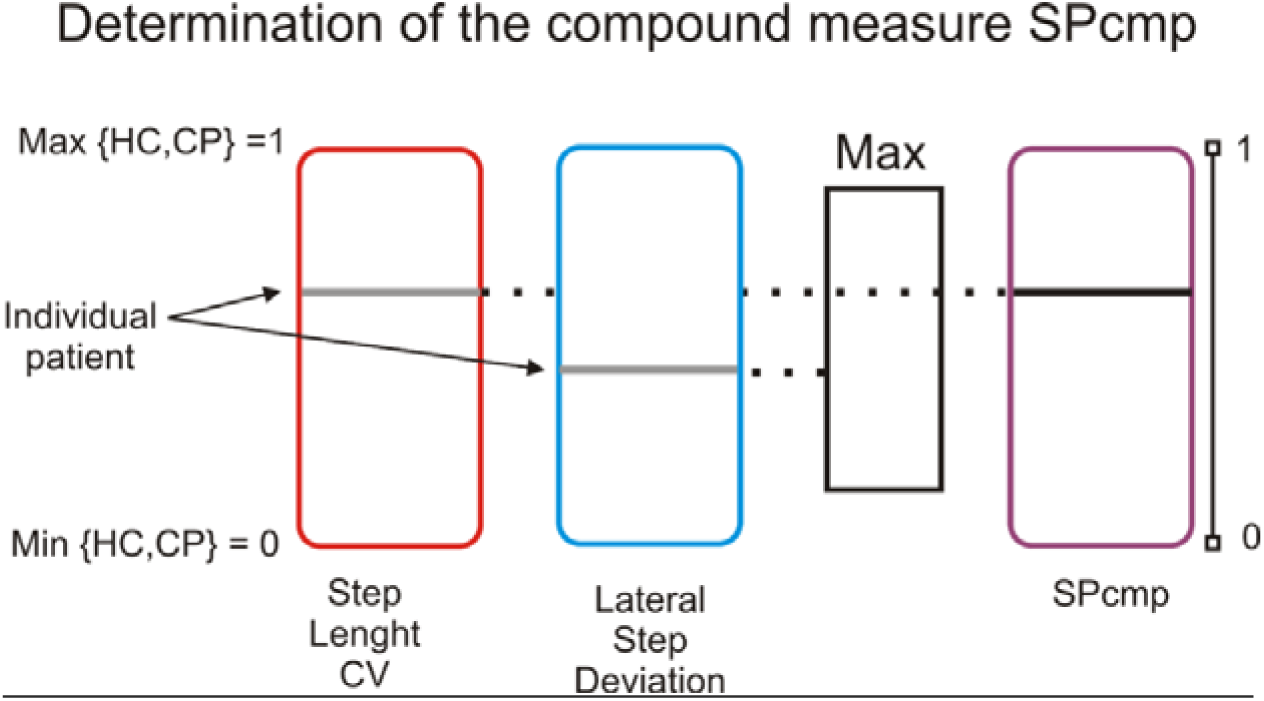
Illustration of the determination of the compound measure SPcmp.

The compound measure combining measures of spatial variability in different dimensions (medio-lateral: *LatStepDev*; aperior-posterior: StrideL_CV_) seems to be effective to capture different components of ataxia-related step-variability. A more detailed analysis on a single-subject level revealed that the component StrideL_CV_ received particular importance for subjects with rather advanced ataxia (Figure 3). These patients performed rather small steps with a small lateral deviation but high variability in the anterior-posterior dimension. Thus, this compound measure can help to capture ataxic gait in different severity stages and using different compensation strategies. Further multi-variate approaches (including the combination of step variability and trunk movement measures) are warranted in the future to capture the variety in real life movements.

**Figure 3.**
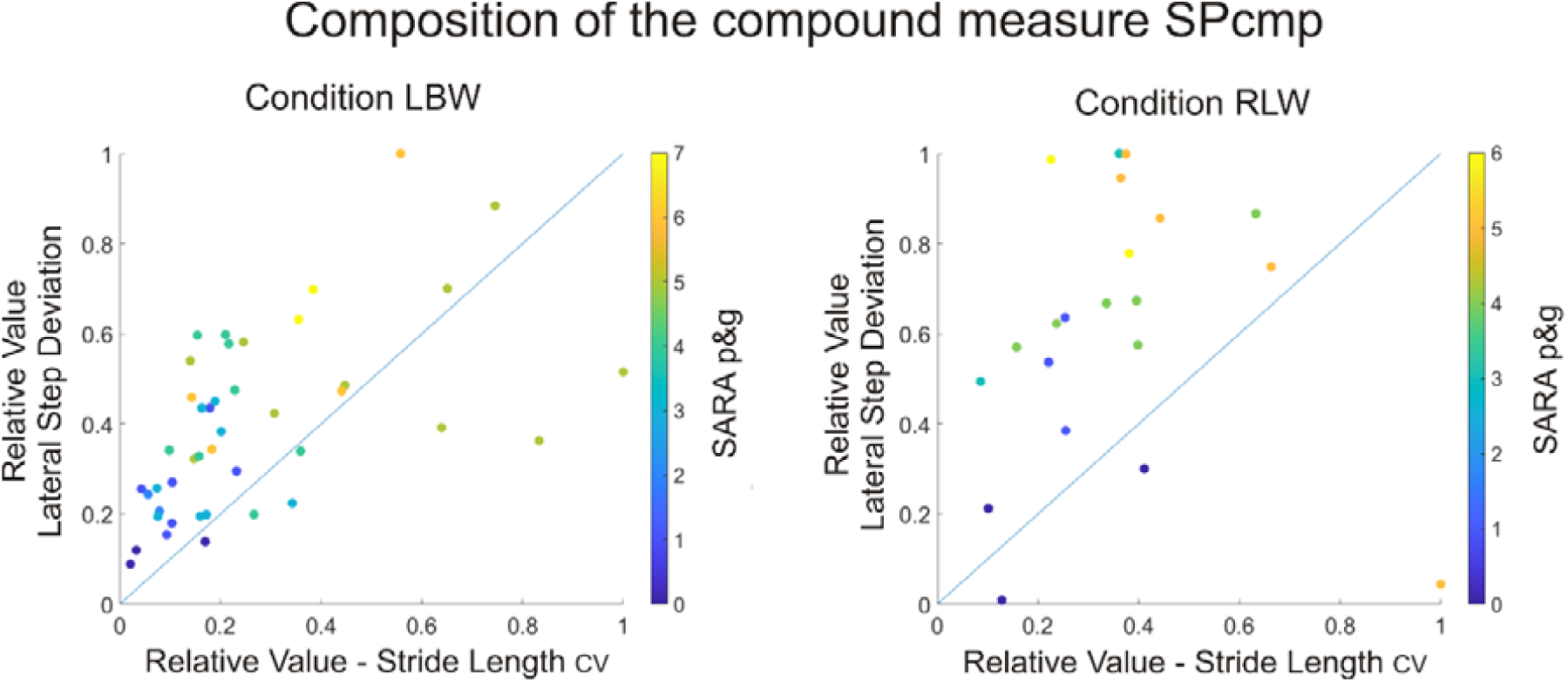
Composition of the compound measure SPcmp for the walking conditions LBW and RLW. Shown are the relative parameter values (see Figure 2) of each patient for the parameters Stride Length CV (x-axis) and Lateral Step Deviation (y-axis). The measure SPcmp is determined by the maximum of the relative values of both measures. The colour coding denotes the severity of gait and posture ataxia as determined by the SARA_p&g_ score.

### Harmonic Ratios

Harmonic ratio (HR) ^35, 36^ of pelvis acceleration was determined to quantify the smoothness of motion. The method quantifies the harmonic content of the acceleration signals (acc) in each direction (HR AP: anterior-posterior, ML medio-lateral, V: vertical) using stride frequency as the fundamental frequency component.

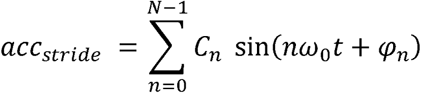

where the C_n_ is the harmonic coefficient, ω_0_ is the stride frequency, and φ_n_ is the phase. A harmonic ratio is calculated by dividing the sum of the amplitudes of the first ten even harmonics by the sum of the amplitudes of the first ten odd harmonics ^35^.

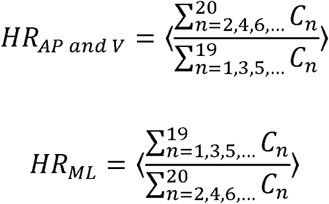

where ⟨ Σ*C*_*n*_ /Σ*C*_*n*_⟩denotes the average ratio over all strides ^52^.

Thus, the HR quantifies the harmonic composition of these accelerations for a given stride, where a higher HR is interpreted as greater walking smoothness.

## Supplement S2 – Results

**Table 1.**
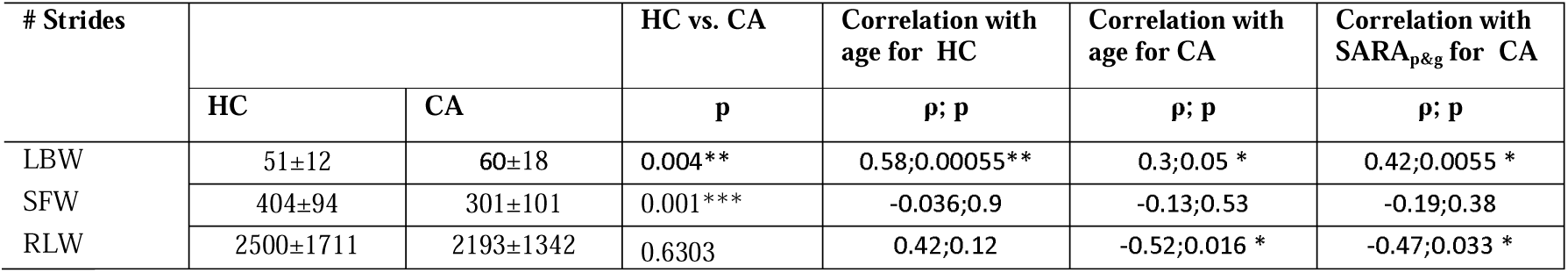
Number of analysed strides for the different walking conditions. Relationships to age and SARA_p&g_ were determined by Spearman correlation.

**Table 2.** Gait measures during the constrained walking condition LBW for the subgroup LBW_RLW-subgroup_ of cerebellar patients that participated also in the real life condition RLW. Given are mean values and standard deviations. HC, healthy controls; SARA_p&g_= SARA posture and gait subscore. (*= p<0.05, **= p<0.007 Bonferroni-corrected, ***= p<0.001).

| Gait measure<br>(in LBW) | HC vs. LBW<sub>RLW-subgroup</sub> |  | Correlation with SARA<sub>p&g</sub><br>for LBW<sub>RLW-subgroup</sub> |  |
| --- | --- | --- | --- | --- |
|  | HC | LBW<sub>RLW-subgroup</sub> | p | p |
| StrideL<sub>CV</sub> | 0.021±0.005 | 0.037±0.017 | 0.00029*** | 0.014* |
| StrideT<sub>CV</sub> | 0.015±0.006 | 0.028±0.01 | <0.0001*** | 0.0011** |
| LatStepDev | 0.019±0.0038 | 0.037±0.01 | <0.0001*** | 0.0003*** |
| SPcmp | 0.12±0.05 | 0.38±0.18 | <0.0001*** | 0.00014*** |
| Harmonic Ratio AP | 3.6±0.95 | 2.7±0.9 | 0.0047** | 0.053 |
| Harmonic Ratio ML | 2.5±0.63 | 2±0.55 | 0.0064** | 0.18 |
| Harmonic Ratio V | 3.3±0.85 | 2.7±0.7 | 0.023* | 0.033* |

**Table 3.** Effect sizes for the group comparison HC vs. CA for the different walking conditions LBW, SFW and RLW. Effects sizes were determined using cohen’s *d*.

| Effects sizes *d* | gait conditions |  |  |
| --- | --- | --- | --- |
|  | LBW | SFW | RLW |
| StrideL<sub>CV</sub> | 3.6 | 0.71 | 0.86 |
| StrideT<sub>CV</sub> | 2.8 | 0.82 | 0.93 |
| LatStepDev | 4.5 | 1.1 | 1.6 |
| SPcmp | 4.4 | 1.8 | 2.6 |
| Harmonic Ratio AP | -1 | -1.4 | -1.4 |
| Harmonic Ratio ML | -0.98 | -1.4 | -1.5 |
| Harmonic Ratio V | -0.95 | -1.3 | -1.4 |

**Table 4.** Accuracy measure quantifying the *classification accuracy* between groups HC vs. CA for the different walking conditions LBW, SFW and RLW. Accuracy was determined by ROC (Receiver operating characteristic).

| HC vs. CA | gait conditions |  |  |
| --- | --- | --- | --- |
| ROC accuracy | LBW | SFW | RLW |
| StrideL<sub>CV</sub> | 0.81 | 0.82 | 0.75 |
| StrideT<sub>CV</sub> | 0.86 | 0.79 | 0.72 |
| LatStepDev | 0.91 | 0.85 | 0.86 |
| SPcmp | 0.91 | 0.85 | 0.86 |
| Harmonic Ratio AP | 0.58 | 0.62 | 0.58 |
| Harmonic Ratio ML | 0.58 | 0.62 | 0.58 |
| Harmonic Ratio V | 0.59 | 0.62 | 0.58 |

**Table 5.** Correlation of gait measures between different walking conditions in cerebellar patients. Effect sizes are given using Spearman’s ρ.

| Correlation of<br>condition LBW | with condition SFW |  | with condition RLW |  | Correlation of<br>condition SFW | with condition RLW |  |
| --- | --- | --- | --- | --- | --- | --- | --- |
| | $\rho$ | p | $\rho$ | p | | $\rho$ | p |
| StrideL<sub>CV</sub> | 0.57 | 0.011* | 0.66 | 0.0019** | StrideL<sub>CV</sub> | 0.63 | 0.0034** |
| StrideT<sub>CV</sub> | 0.64 | 0.0028* | 0.42 | 0.068 | StrideT<sub>CV</sub> | 0.44 | 0.053 |
| LatStepDev | 0.84 | <0.0001*** | 0.74 | 0.0003*** | LatStepDev (%) | 0.66 | 0.0021** |
| SPcmp | 0.85 | <0.0001*** | 0.83 | <0.0001*** | SPcmp | 0.8 | <0.0001*** |
| Harmonic Ratio AP | 0.8 | <0.0001*** | 0.46 | 0.044* | Harmonic Ratio AP | 0.64 | 0.003** |
| Harmonic Ratio ML | 0.81 | <0.0001*** | 0.57 | 0.01* | Harmonic Ratio ML | 0.64 | 0.0032* |
| Harmonic Ratio V | 0.82 | <0.0001*** | 0.15 | 0.53 | Harmonic Ratio V | 0.31 | 0.19 |

**Table 6.** Correlations of gait measures between different walking conditions in healthy controls. Effect sizes are given using Spearman’s ρ.

| Correlation of<br>condition LBW | with condition SFW |  | with condition RLW |  | Correlation of<br>condition SFW | with condition RLW |  |
| --- | --- | --- | --- | --- | --- | --- | --- |
| | $\rho$ | p | $\rho$ | p | | $\rho$ | p |
| StrideL<sub>CV</sub> | 0.06 | 0.85 | 0.32 | 0.29 | StrideL<sub>CV</sub> | 0.74 | 0.0058** |
| StrideT<sub>CV</sub> | 0.38 | 0.2 | 0.18 | 0.55 | StrideT<sub>CV</sub> | 0.54 | 0.61 |
| LatStepDev | 0.71 | 0.0081* | 0.46 | 0.11 | LatStepDev (%) | 0.55 | 0.055 |
| SPcmp | 0.62 | 0.029* | 0.41 | 0.16 | SPcmp | 0.47 | 0.11 |
| Harmonic Ratio AP | 0.81 | 0.0014** | 0.32 | 0.29 | Harmonic Ratio AP | 0.42 | 0.15 |
| Harmonic Ratio ML | 0.73 | 0.0069** | 0.24 | 0.43 | Harmonic Ratio ML | 0.51 | 0.078 |
| Harmonic Ratio V | 0.77 | 0.0029** | 0.75 | 0.0044** | Harmonic Ratio V | 0.64 | 0.021 |

**Table 7.** Correlations between trunk and step measures for the different walking conditions determined by Spearman correlation. Effect sizes are given using Spearman’s ρ.

|  | Correlation of harmonic ratio AP in condition | with StrideL<sub>CV</sub> |  | with StrideT<sub>CV</sub> |  | with LatStepDev |  | with SPcmp |  |
| --- | --- | --- | --- | --- | --- | --- | --- | --- | --- |
| | | $\rho$ | p | $\rho$ | p | $\rho$ | p | $\rho$ | p |
| cerebellar patients | LBW | -0.55 | 0.0002*** | -0.59 | 0.00013*** | -0.46 | 0.0021** | -0.55 | 0.00014*** |
|  | SFW | -0.38 | 0.06 | -0.46 | 0.026 * | -0.45 | 0.03* | -0.48 | 0.018* |
|  | RLW | -0.76 | 0.0006*** | -0.67 | 0.0011** | -0.42 | 0.05 | -0.61 | 0.0036 ** |
| healthy controls | LBW | -0.59 | 0.00052** | -0.29 | 0.11 | -0.28 | 0.13 | -0.37 | 0.044* |
|  | SFW | -0.06 | 0.82 | -0.14 | 0.61 | -0.29 | 0.29 | -0.33 | 0.23 |
|  | RLW | -0.54 | 0.04* | -0.27 | 0.33 | -0.47 | 0.076 | -0.41 | 0.13 |
|  | Correlation of harmonic ratio ML in condition | with StrideL<sub>CV</sub> |  | with StrideT<sub>CV</sub> |  | with LatStepDev |  | with SPcmp |  |
| cerebellar patients | LBW | -0.31 | 0.046* | -0.26 | 0.09 | -0.37 | 0.015* | -0.37 | 0.016 * |
|  | SFW | -0.65 | 0.0007** | -0.36 | 0.08 | -0.61 | 0.002** | -0.62 | 0.0013** |
|  | RLW | -0.4 | 0.072 | -0.58 | 0.007* | -0.5 | 0.02* | -0.54 | 0.011* |
| healthy controls | LBW | -0.39 | 0.032 | -0.08 | 0.65 | 0.16 | 0.38 | 0.037 | 0.84 |
|  | SFW | -0.21 | 0.46 | -0.32 | 0.24 | -0.27 | 0.32 | -0.3 | 0.28 |
|  | RLW | -0.16 | 0.57 | -0.2 | 0.47 | -0.25 | 0.36 | 0.08 | 0.77 |
|  | Correlation of harmonic ratio V in condition | with StrideL<sub>CV</sub> |  | with StrideT<sub>CV</sub> |  | with LatStepDev |  | with SPcmp |  |
| cerebellar patients | LBW | -0.63 | <0.0001*** | -0.65 | 0.0001*** | -0.52 | 0.0003*** | -0.6 | 0.0001*** |
|  | SFW | -0.54 | 0.007 * | -0.42 | 0.04* | -0.57 | 0.004** | -0.59 | 0.002 ** |
|  | RLW | -0.67 | 0.0012** | 0.59 | 0.0054** | -0.43 | 0.051 | -0.59 | 0.005 ** |
| healthy controls | LBW | -0.23 | 0.21 | -0.09 | 0.62 | -0.14 | 0.46 | -0.2 | 0.29 |
|  | SFW | -0.11 | 0.69 | -0.24 | 0.38 | -0.42 | 0.12 | -0.46 | 0.083 |
|  | RLW | -0.35 | 0.21 | -0.2 | 0.47 | -0.44 | 0.11 | -0.26 | 0.34 |

